# Decline of Common Toad populations in Flanders is not linked to surrounding landscape

**DOI:** 10.64898/2026.01.07.698094

**Authors:** Ellen Blomme, Henri Rommel, Femke Batsleer, Lieven Clement, Dominique Verbelen, An Martel, Siska Croubels, Frank Pasmans, Dries Bonte

**Affiliations:** Wildlife Health Ghent, Faculty of Veterinary Medicine, Ghent University, Salisburylaan 133, Merelbeke, Belgium; Terrestrial Ecology Unit, Department of Biology, Faculty of Science, Ghent University, K.L. Ledeganckstraat 35, Ghent, Belgium; Mathematics, Computer science and Statistics, Faculty of Science, Ghent University, Krijgslaan 281, Ghent, Belgium; Natuurpunt Studie, Coxiestraat 11, Mechelen, Belgium; Laboratory of Pharmacology and Toxicology, Faculty of Veterinary Medicine, Ghent University, Salisburylaan 133, Merelbeke, Belgium

**Keywords:** amphibian decline, Anura, common species, land use, landscape heterogeneity, trend analysis

## Abstract

**Aim:** We aim to quantify population trends of Common Toad in Flanders and assess to which extent population trends depend on general landscape variables (land use identity, structure and change).

**Location:** Flanders, Belgium

**Methods:** Using standardized time series obtained from citizen science and spanning over four decades, we used an end-begin contrast to get a trend value for 234 populations. Next, we developed a testing strategy to find associations between these trend values and the surrounding landscape

**Results:** Across the study region, 40% of the populations have declined significantly, while only 10% show an increase. Declines were not associated with landscape characteristics in our study area, which is one of the most fragmented regions in Europe.

**Main conclusions:** We discuss how the research region, spatial and temporal resolution, as well as the generalist nature of the species, may explain these findings. Our study suggests limited effects of general landscape characteristics on the decline of Common Toad populations, indicating that other major drivers are likely responsible.

## Introduction

Anthropogenic impacts are the primary drivers of the current global biodiversity loss (Gaston and Fuller 2008; Dirzo et al. 2014; Cowie et al. 2022). Among vertebrates, amphibians are experiencing some of the steepest population declines worldwide (Stuart et al. 2004). In Europe, 59% of the amphibian taxa are declining of which 23% are threatened with extinction (Temple and Cox 2009; López-de Sancha et al. 2025). While biodiversity loss has spurred research and conservation of rare and threatened species (Gaston and Fuller 2008), common and more widespread species often receive disproportionately less attention. Yet, even relatively small declines in numerically dominant species can disrupt ecosystem structure and functioning. Importantly, for more widespread species—in contrast with persistently rare species—conservation actions still hold a large potential to reverse or mitigate (Langhammer et al. 2024).

The Common Toad (*Bufo bufo*) exemplifies this concern. As one the most widespread European amphibians, it plays a crucial role in energy transfer between aquatic and terrestrial systems (Gibbons et al. 2006). The species is a mesopredator that feeds on a broad array of invertebrate species, while also serving as an important prey for predators, such as herons, corvids, rats and polecats (Mertens and Snep 2009). The IUCN currently lists the Common Toad as Least Concern because of its wide distribution, presumed large populations and unlikeliness to be declining fast enough (IUCN SSC Amphibian Specialist Group 2023). However, despite stable distribution patterns, significant and widespread declines in abundance have been reported across several European countries, including the UK (Carrier and Beebee 2003; Petrovan et al. 2025), Italy (Bonardi et al. 2011), Switzerland (Petrovan et al. 2025) and the Netherlands (Ravon 2021). These trends underscore the need to reassess conservation priorities and monitoring strategies for species traditionally considered secure. Declines in Common Toad are likely driven by multiple interacting stressors operating across multiple spatial scales, affecting reproduction and survival throughout the species’ life cycle (Nolan et al. 2023). Key drivers of this decline include habitat loss (e.g., loss of ponds and small landscape elements), connectivity loss (including migratory connectivity between breeding ponds and adult terrestrial habitats), climate change, loss of genetic diversity, diseases and land use (e.g. agricultural cultivation practices and pesticide use). These local scale processes primarily affect the aquatic phase, influencing tadpole development, habitat and survival. In contrast, adult toads have large home ranges, often extending up to 3 km from their breeding ponds. These larger-scale effects are likely to influence adult body condition and survival (Biek et al. 2002; Harper et al. 2008; Bonardi et al. 2011; Petrovan and Schmidt 2016).

Landscapes typically comprise a mosaic of land use types, each with distinct effects on populations. This is referred to as identity effects of the landscape (Landler et al. 2023). Land use categories, such as woodlands, agricultural land and urbanized areas differ in habitat suitability, resource availability and environmental conditions that directly impact amphibians (Hartel et al. 2008; Salazar et al. 2016). Beyond these identity effects, the structure and organization of land use types within landscapes also play a vital role. Landscapes may be homogenously organised when one terrestrial habitat dominates or heterogeneously organised when a mix of habitats coexist. Landscape heterogeneity can further be subdivided into compositional heterogeneity (variation of land-cover types) and configurational heterogeneity (spatial arrangement of land-cover components). Compositional heterogeneity introduces a variety of environmental conditions, while configurational heterogeneity influences ecological processes within and between habitat patches (Li and Reynolds 1995; Tonetti et al. 2023). These two dimensions are often tightly interlinked as mixed landscapes often contain several small patches resulting in more fragmentation (Fletcher 2005). Fragmentation can alter metapopulation dynamics through connectivity loss and amplify edge effects (Saunders et al. 1991; Schlaepfer and Gavin 2001).

For a common species like the Common Toad, continuous habitat areas may reduce fragmentation and edge-related stressors, potentially supporting better population health. However, heterogenous landscapes may offer greater resource availability, as the main food resources, invertebrates, have generally higher biomasses and biodiversity in such landscapes (Sinclair et al. 2024). In summary, both the identity and the compositional and configurational heterogeneity of landscapes influence amphibian population trends across spatial scales.

We used a large citizen science dataset of toad patrols across Flanders to address the following research questions 1) what are the population trends of Common Toads and 2) how do landscape variables (land use identity, structure and change) affect changes in Common Toad population trends? This dataset provides a unique opportunity to assess long-term patterns in abundance and distribution, while linking observed changes to landscape-level processes across spatial scales.

## Methods

### Trend analyses

We assessed the extent of decline in the research area, Flanders (Belgium), for the Common Toad (*Bufo bufo*). Given the strong annual variability in toad population sizes, sufficiently long time series are essential to reliably estimate the population trend. To this end, we used the data collected during the spring migration of toads in Flanders by volunteers of nature conservation organisation “Natuurpunt Studie” since 1981 until 2022, as dataset previously used in Blomme et al. (2025). Data cleaning was performed using the ‘janitor’ package (Firke 2023). We ensured standardized names and unique names for each location and assumed that each sampling location represented a distinct population. To distinguish trends from demographic fluctuations, only populations that had at least six observations of data were included (Meyer et al. 1998; Green 2003). Most time series extended at least until 2014, with only 14 exceptions (supplementary Table S1). The longest time series, spanning from 1988 to 2021 included 30 observations. In total, we analysed the trend for 234 out of the initial 640 populations (supplementary Table S1). All statistical analyses were performed in R v4.3.2 (R Core Team 2023), data manipulation and visualisation was done with the R-package collection ‘tidyverse’ (Wickham et al. 2019) and geospatial operations were performed with a combination of R and QGIS (QGIS Development Team 2024).

As the response variable, we used the annual sum of all recorded individuals per population, including both roadkills and migrating toad records. We modelled the counts using the negative binomial distribution to account for overdispersion regarding the Poisson distribution. Although serial correlation commonly occurs in ecological time series, autocorrelation function (ACF) and partial autocorrelation function (PACF) did not indicate any autocorrelation (Hyndman and Athanasopoulos 2021). In 60% of the populations, gaps in the time series were present.

Population dynamics were often nonlinear, necessitating the use of flexible modelling approaches such as Generalized Additive Models (GAMs) (Hastie et al. 2017). Accordingly, we modelled all the populations with sufficient long time series as GAM (at least 10 observations), while all short time series were fitted with GLM (6, - 9 observations), with the packages ‘MASS’ (Venables and Ripley 2002), ‘boot’ (Canty and Ripley 2022) and ‘mgcv’ (Wood 2003, 2004, 2011, 2017; Wood et al. 2016). Trend inference for GAMs was based on omnibus tests followed by post-hoc tests with multiple testing correction using the stage-wise analysis from the ‘stageR’ package (Van den Berge et al. 2017). The omnibus test evaluates the overall significance of the smoother, using a likelihood ratio test. In the post-hoc analysis we assessed two contrasts. The Begin vs End contrast reflects the overall trend and consists of the difference between the fit at the last timepoint and the first timepoint, divided by the number of years (to correct for time span). The Average Slope approach evaluates the rate of change over time and captures both the magnitude and direction of the change and was calculated by taking the average of the first derivative of the smoother. Both contrasts were highly positively correlated (Spearman correlation, r_s_(109) = 0.93, p < 0.001). From a biological perspective, the begin-end contrast was considered more meaningful, even when intermediate fluctuations occur. Therefore, trend classification was based on this contrast.

Each population was labelled after correcting for multiple testing. When the omnibus test did not reveal a significant trend (p_adj_ > 0.05), populations were labelled ‘non-significant’. If a trend was detected, further conclusions were based on the post-hoc test. Populations were labelled ‘increasing’ or ‘decreasing’ if the trend value was positive or negative respectively and the post-hoc test passed the 5% overall FDR threshold (Van den Berge et al. 2017). If only the omnibus null hypothesis was rejected, the population was labelled ‘flexible’, indicating non-monotonic dynamics.

### Landscape analyses

To characterize the landscape context around toad patrol locations, we used the 2018 map ‘Land cover and Land use Flanders’ (Bodembedekking & Bodemgebruik Vlaanderen, BBK; geopunt.be; Vlaanderen 2021). This raster-based map, produced by the Flemish government, has a resolution of 1 m^2^ and is based on remote sensing and field observations. We reclassified the original 14 into 10 land use components, excluding four categories indicating canopy hanging over water or roads, which were deemed biologically irrelevant for Common Toad biology. The resulting components included: buildings, roads, covered, uncovered, grass & shrubs, railways, water, woodland, arable land and agricultural grasslands. An 11th category (biological valuable grasslands) was added from the Biological Valuation Map (Biologische waarderingskaart; De Saeger et al. 2023, BWK; geopunt.be) to separate ‘gardens & roadsides’ from ‘(semi-)natural grasslands’ (Table S2).

To quantify the landscape context and structure at different scales, we generated three circular buffers around each of the toad migration sampling locations: 100 m, 500 m and 1000 m using QGIS (QGIS Development Team 2024). These scales were chosen to capture both local and broader landscape effects. The smallest buffer reflects immediate habitat conditions, while the largest buffer accounts for landscape-level influences on adult toads, which may range up to 3 km from breeding ponds. The intermediate scale was selected based on known patterns of toad movement and hibernation distances prior to spring migration (Sinsch 1988; Salazar et al. 2016).

We calculated the proportions of land use components within each buffer zone (100 m, 500 m, and 1000 m) for every toad patrol location using the ‘terra’ (Hijmans et al. 2024), ‘raster’ (Hijmans et al. 2023) and ‘sf’ (Pebesma et al. 2024) packages. To explore landscape variation across populations, we performed a Principal Component Analysis (PCA) with z-transformed proportions of the 11 initial land use components using the ‘vegan’ package (Oksanen et al. 2022). Furthermore, correlograms were constructed to understand correlation among land use components. Positively correlated and biologically similar components were merged into broader categories (Table S2), resulting in six remaining land use components (agricultural land, woodland, water, urbanized area, biological valuable grasslands and railways). Furthermore, we calculated three metrics that represent the structure and heterogeneity of the landscape: the Shannon Diversity Index based on the six identity components as a measure for compositional heterogeneity, the number of patches and edge length as measures for configurational heterogeneity. Note we explored more metrics, however a lot of landscape metrics have a large redundancy among them. To assess land use change, we extracted land use component proportions from the 2012 and 2021 versions of the BBK map. We then calculated the Bray-Curtis distance between the land use in 2012 and the land use in 2021 from a single site and used this as a proxy for landscape transformation over time.

To investigate how landscape variables influence the trend value of Common Toad (based on the begin-end contrast), we constructed a linear model using the six land use identities, three structural variables and one variable for land use change. However, due to the compositional nature of the dataset, where land use proportions sum to approximately one, aliasing occurs among identity components, resulting in high collinearity. Indeed, the land use proportion for the last class can be replaced by one minus the sum of the proportions of the remaining classes. Specifically, we modified the linear model by leaving out the predictor agricultural land. Note that the interpretation of the intercept then changes to average trend value when the area around the Common Toad population consists 100% of agricultural land and the slope terms for the remaining classes are corrected for the fact that an increasing use implies a reduction in agricultural land (supplementary information S3).

We first assessed whether interactions among the land use identities should be included in the model. Specifically, we compared a model with main effects for each land use identity, structure and change with a model including these main effects and all two-way interactions involving land use identities using an F-test. If no significant difference in fit was observed, we retained the main effects model, and we conducted an omnibus test to assess the association of all land use predictors with the Common Toad population trends, simultaneously. Again using an F-test to compare the selected model to an intercept only null model. If either the omnibus test for the interaction or the omnibus test for the main effects were significant, we conducted post-hoc tests to pinpoint significant associations.

## Results

### Trend analyses

The 234 migration sampling locations scattered across Flanders, for which a trend was calculated, have an average time series length of 10.24 years and collectively accounted for more than 1.7 million migrating toads. Of these, 111 populations had sufficiently long time series (≥10 observations) to be evaluated using GAM with the omnibus and post-hoc tests (Fig. 1a-b). Among these, 33 populations showed no significant trend. Among the 78 populations that initially showed a statistically significant trend based on the omnibus test, 16 populations did not yield statistically significant results in the post-hoc test. Visual inspection of these trends suggests these exhibited a flexible trend (yellow in Fig. 1b), with almost the same value for start and end point but variability in the intermediate period. Ultimately, 62 out of 111 populations displayed statistically significant trends in both the omnibus and post-hoc tests. From these, only 12 populations exhibited a significant increase in population, while 50 populations experienced a significant decrease.

**Fig. 1.**
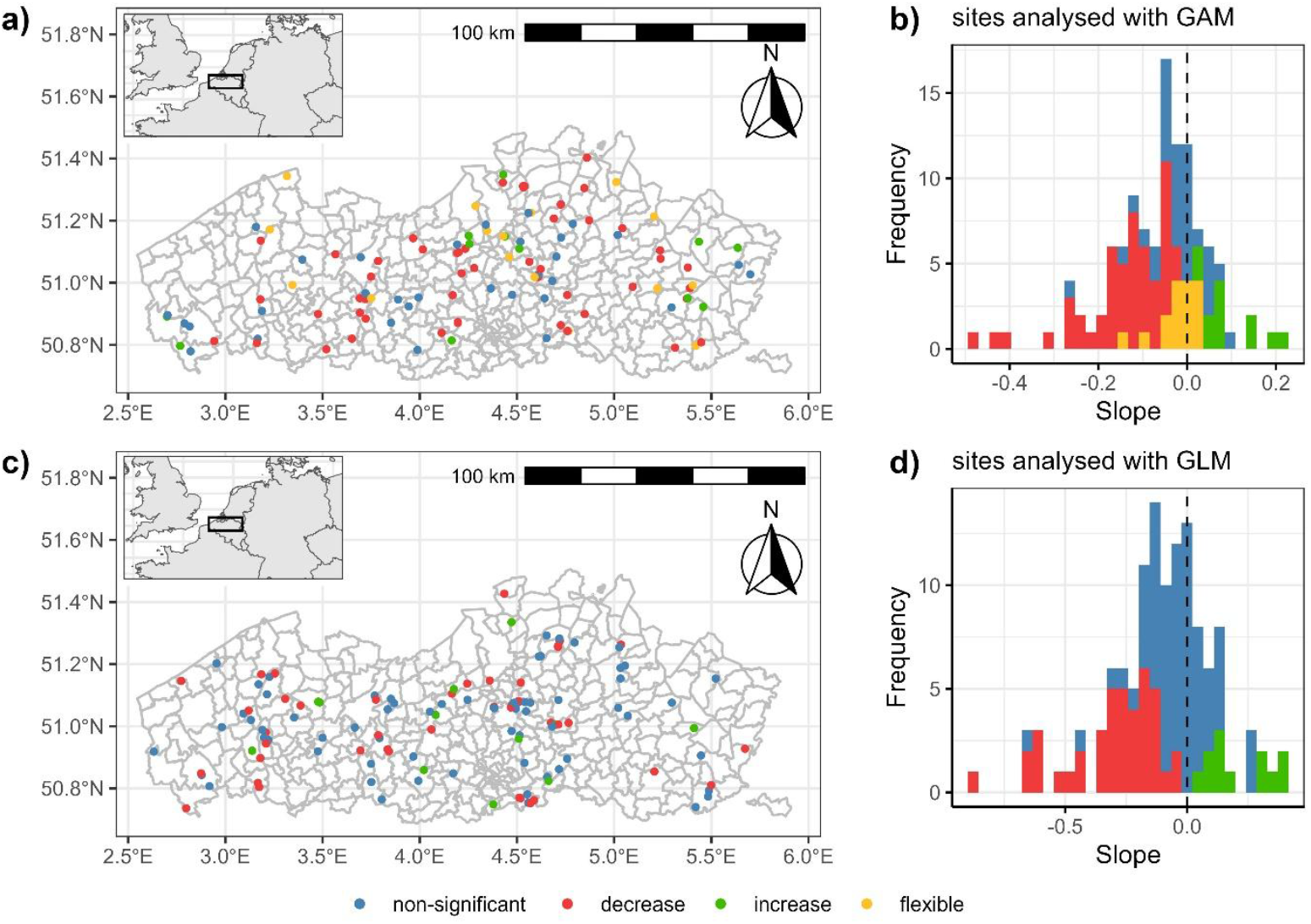
Visualisation of the different sample locations in Flanders, with the colours representing the status of the trends. a) the distribution of the populations with a sufficiently long time series (at least 10 observations; N=111), modelled using GAM with an inset map that provides the location of Flanders within northwestern Europe and b) histograms of the slopes of the populations estimated with GAM; c) the distribution of the populations with shorter time series (6-9 observations; N=123), modelled using GLM and c) histograms of the slopes of the populations estimated with GLM

The remaining 123 population sampling locations, which had shorter time series, were evaluated using GLM (Fig. 1c-d). Over half of these populations displayed no significant trend after correction for multiple testing. Among the 53 populations with short time series exhibiting significant trends, only 12 populations demonstrated evidence of population increase, while 41 populations showed a decline in population size. In total we found 91 out of 234 populations that had a significant decline in population, of which 36 completely disappeared.

### Landscape analysis

A PCA ordination (Fig. 2, supplementary Table S4 and Fig. S5) was used to explore landscape composition variation among the Common Toad populations. These results confirm that this generalist species occurs across a variety of different landscapes. At the 500 m scale (Fig. 2), and similar for the two other scales (supplementary Fig. S5), PC1 shows a gradient from more urban land use components, such as buildings and roads, to more rural components such as woodlands, semi-natural grasslands (biological valuable grass & shrubs) and arable land. The PC2 axis captured a gradient from sparser, less-covering vegetation to denser vegetation such as woodlands. Urban land use components (buildings, roads, covered, uncovered, grass & shrubs) are consistently positively correlated. This is already apparent on the 100 m scale and is enhanced in the 500 m (Fig. 2) and 1000 m scales (supplementary Fig. S5).

**Fig. 2.**
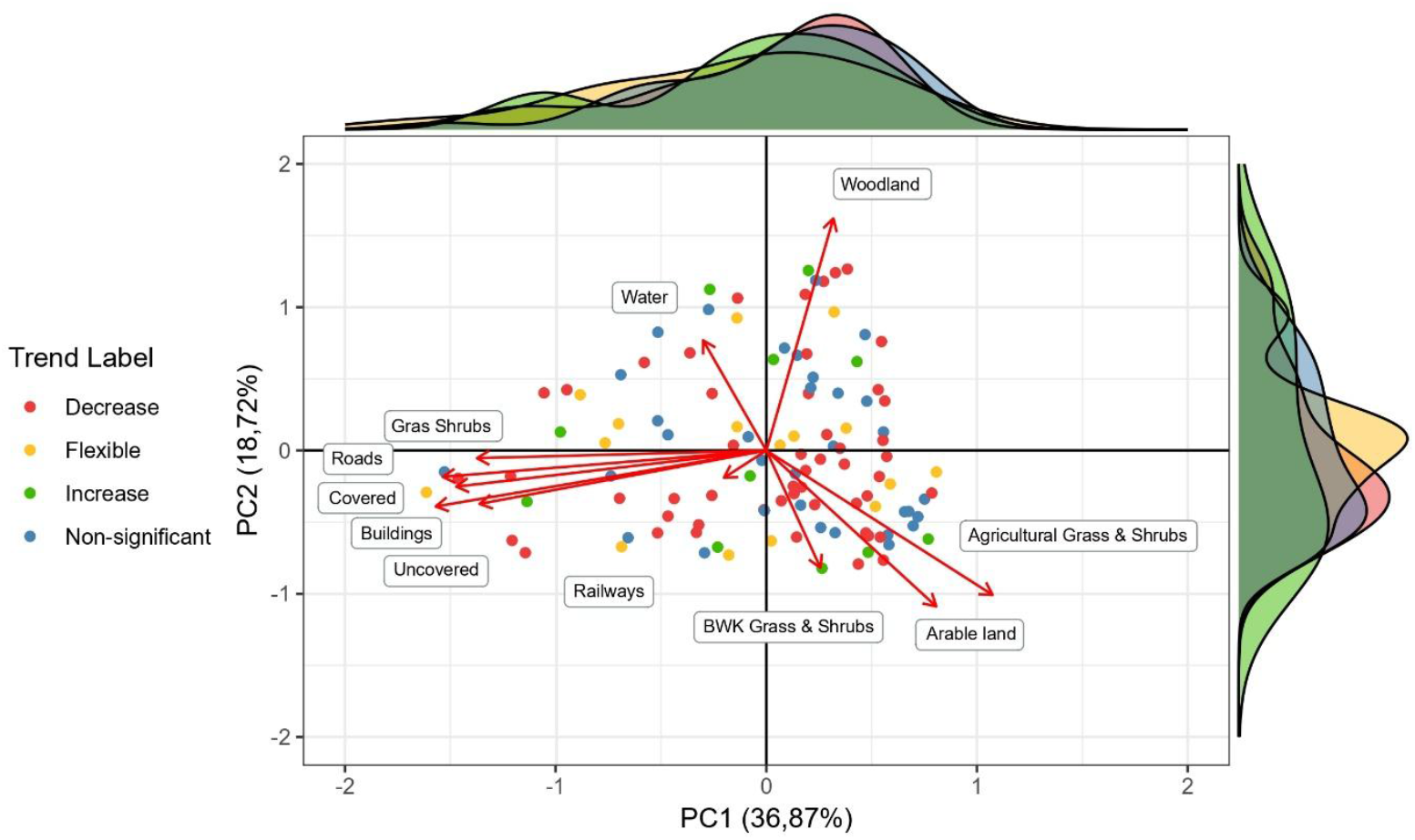
PCA ordination biplot for the 500 m buffer of the landscape composition proportions. Different colours detail different labels of toad population trends. Marginal density graphs describe the density of different populations corresponding to different trend labels. Similar plots for the 100 m and 1000 m buffer scales are in supplementary Fig. S4

Based on the earlier correlations between land use identities, we move from 11 to 6 components to construct these linear models. We first evaluated whether interactions between land use identities should be included in the model. According to the omnibus tests none of the two-way interactions amongst land use identities were statistically significant at any spatial scale: 100 m (F_9,101_ = 1.296, p = 0.2225), 500 m (F_9,101_ = 1.0953, p = 0.3730) and 1000 m (F_9,101_ = 1.4008, p = 0.1654). Given these results, we used the main effects models to further study the associations between the Common Toad trends and the landscape characteristics. Next, we used an omnibus test to assess whether at least one of the landscape characteristics is associated with the trend. Across all examined spatial scales, none of the associations were statistically significant (Table 1) and cannot be used to explain the variation in trends.

**Table 1:** Overview of F-tests to compare the full models without interactions to the null model. Given columns are buffer scale, F-value of a model, the degrees of freedom (Df) and the associated p-value.

| Buffers | F-value | Df | p-value |
| --- | --- | --- | --- |
| 100 m | 1.735 | 101 | 0.09057 |
| 500 m | 1.208 | 101 | 0.2983 |
| 1000m | 1.063 | 101 | 0.3964 |

## Discussion

As observed in other European countries, we found local declines of Common Toad abundances across Flanders (Carrier and Beebee 2003; Bonardi et al. 2011; Petrovan et al. 2025). Nearly 39% of the populations showed decreasing trends, while only 10% of the local populations increased in numbers. Alarmingly, of these declining populations, 40% reached such low densities that toad patrol actions have been discontinued. Compared to other countries, we found less declining populations than Italy, who found 90% of the investigated populations to decline (Bonardi et al. 2011), but more than the 30% decline in the UK (Carrier and Beebee 2003). Although these declines are likely driven by multiple interacting stressors, we focused on the putative association between general surrounding landscape and these trends, as a first and important step to uncover underlying mechanisms. To achieve this, we developed a testing strategy to tackle unit-sum proportion data. However, our analysis revealed that land use identity, structure and change did not significantly explain variation in population trends, regardless of spatial scale. Flanders (Belgium) is one of the most fragmented areas in Europe characterised by a mosaic of different land uses (EEA 2011). As a result, homogenous landscapes in the strict sense do not occur in our study region. Toad patrols are also founded in areas as a conservation measure against road mortality in amphibians (Schmidt et al. 2008). This means that at larger spatial scales these locations are not randomised and will always include similar anthropogenic components (Petrovan et al. 2025). Although our dataset included a high number of locations, the landscape matrix surrounding the ponds is relatively similar, particularly on a larger scale. This limited variability on both land use identity and heterogeneity (compositional and configurational) may have reduced our ability to detect associations between population trends and landscape variables.

We did not find a general association between Common Toads and surrounding landscape characteristics, but localised and specific combinations of landscape characteristics may still exert biological relevant impact. There might still be an impact of specific landscape characteristics on other specific population metrics, individual traits or behaviour. In Common Toad, Bettencourt-Amarante et al. (2025) showed consequences of land use identity on morphological traits, which can lead to lower individual health (Guillot et al. 2016). In fragmented landscapes, juvenile toads are highly motivated to move and explore, but are more vulnerable to desiccation, exhaustion or predation due to their sensitivity to water (Janin et al. 2012). Furthermore, adult toads have to cross more landscape boundaries, which might result in higher dispersal costs (Schultz et al. 2012; Potts et al. 2016). Salazar et al. (2016) mapped the relative occurrence of Common Toads and have shown that this increases with the distance to water bodies and wooded habitats. Toads are also known to occur less in farmland itself and will often prefer more hospitable landscape features within agricultural areas (Taylor et al. 2024). Similarly, densities of Common Toad are often higher in wooded habitat (Latham 1997). In Wood Frog (*Lithobates sylvaticus*), it has been found that fecundity responds differently to the landscape context than abundance or occurrence (Moraga et al. 2019). These examples highlight that landscape impacts on toads might be shaped by specific ecological interactions, which might not be detected with our study looking into general landscape-context effects on the overall declines in populations.

In addition to land use identity and heterogeneity, we also did not find any associations with land use change. Our analyses covered a nine-year time interval using high resolution thematic maps. However, the limited temporal span may have constrained the detection of meaningful changes, either due to minimal landscape transformation or the presence of time-lag effects in ecological responses. Another potential limitation might be that changes in small landscape elements are not easily detected by our coarse method, especially since we know that Common Toads benefit from small landscape elements that provide habitat corridors (Taylor et al. 2024). Such restrictions can obscure the detection of a land use change effect. The importance of historic land use has been shown in other studies for the occurrence of three palearctic amphibian species, including the Common Toad (Piha et al. 2007). However, this study focuses on a finer spatial resolution and a larger temporal interval, examining more localized habitat changes. Also, in a *Triturus* species, it has been shown that abundances and assemblages are sensitive to dynamics in landscape corridors and agrarian reform (Arntzen 2025).

As we did not detect general landscape effects on population trends, such large effects of land use might be of minor importance as Common Toads are habitat generalists. Across its range, the Common Toad occupies a diverse set of habitats, both natural environments such as forests or marshes, and anthropogenic areas such as gardens or parks with man-made ponds (Speybroeck et al. 2018). While woodlands often support the highest densities of Common Toads (Latham 1997), studies also report their presence in agricultural areas such as vineyards (Leeb et al. 2020) or urban environments (Budzik et al. 2013; Mazgajska and Mazgajski 2020). This ecological flexibility suggests that Common Toads can persist across diverse land use types. The limited influence of these landscape variables imply that research efforts should not solely focus on general landscapes but prioritize other drivers at local scales such as small landscape elements. The observed variation in population trends may not only be attributable to surrounding landscape, but other drivers might be at play. Such other factors might include climate change, diseases, environmental contaminants and loss of genetic diversity, which all could play a more significant role in structuring population dynamics (Blaustein and Kiesecker 2002). For example, climate change is changing the phenology of toad migration, both timing and duration which can affect species interactions and survival (Blomme et al. 2025). This illustrates the multifactorial cause of decline, which means that we should aim to minimize effects of land use to reduce cumulative pressures in fragmented environments.

In this study, we quantified the extent of the decline of Common Toad populations in Flanders using a large-scale citizen science dataset. We found that about 40% of the populations are declining. We contributed to landscape methodology by developing a testing strategy to find associations between the surrounding landscape and population dynamics, within unit-sum proportion datasets. Nevertheless, we could not link population trends to land use identity, structure or changes at three relevant spatial scales. This may be due to the limited landscape variability in Flanders, a too coarse spatial and/or temporal resolution, or the species’ ecological flexibility. While Common Toads can still be affected by specific landscape characteristics not detected with our general landscape metrics, our results suggest that local land use is not the primary driver of Common Toad declines. Instead, other non-correlated factors such as climate change, disease, contaminants and loss of genetic diversity are likely to have more significant impact on population dynamics (Blaustein and Kiesecker 2002). For conservation, this means that actions should prioritize specific land use characteristics rather than general land use and minimizing cumulative pressures as the decline is most likely driven by multiple factors.

## Acknowledgements

We thank all the volunteers from Natuurpunt that assisted in the toad patrols throughout the years.

## Data Accessibility Statement

The data was deposited in Zenodo (https://doi.org/10.5281/zenodo.18164035). Raw count data are owned by Natuurpunt. This embargo is one of copyright, since the data are collected by volunteers, that remain owner of their data. Due to ethical reasons (privacy of volunteers) the exact coordinates of the patrols cannot be shared.

