## Supplementary Material for "Decline of Common Toad populations in Flanders is not linked to surrounding landscape"

* Shared first authors

**Supplementary material**

Supplementary Table S1:

This Table gives the output for each population with a time series longer than 5 years. For each toad patrol, there is a unique location number, street name and province. The adjusted p values from the omnibus test and the post-hoc test are shown and together with the trend value, the status of the toad population (label; D = decrease, I = increase, F = flexible and NS = non-significant) can be assigned to the population. When the omnibus test is non-significant (omnibus p_adj_ > 0.05), the trend of the population is non-significant. Otherwise there is a trend, which is either flexible (p_adj_ > 0.05), increasing or decreasing, depending on the sign of the trend value. Furthermore, the kind of model that is used to estimate the trend of the population (GAM = Generalized Additive Model; GLM = Generalized Linear Model), amount of observations (# obs) and the timeframe (start-end) within they are observed are given. When the trend is estimated with a GLM, there is no omnibus p-value, which results that these sites cannot have flexible as label.

| **Site id** | **Street name** | **Province** | **Omnibus p_adj_** | **p_adj_** | **Trend value** | **Label** | **Model** | **Start** | **End** | **# obs** |
| --- | --- | --- | --- | --- | --- | --- | --- | --- | --- | --- |
| 5 | Abdij Van Park | Vlaams Brabant | 1.66E-02 | 1.11E-03 | -0.27 | D | GAM | 2010 | 2021 | 13 |
| 10 | Aertschouw (Opoeteren) | Limburg | 9.19E-01 |  | 0.01 | NS | GAM | 2011 | 2022 | 12 |
| 15 | Appeldijkstraat (Weert) | Antwerpen | 2.61E-03 | 2.81E-02 | -0.06 | D | GAM | 1994 | 2007 | 11 |
| 16 | Arendsnest | Antwerpen | 2.90E-05 | 2.88E-10 | 0.15 | I | GAM | 2009 | 2022 | 12 |
| 25 | Ballewijerweg | Limburg | 6.87E-03 | 1.91E-03 | -0.05 | D | GAM | 1988 | 2022 | 17 |
| 27 | Bankelindeweg | West-Vlaanderen | 2.18E-06 | 2.45E-02 | 0.03 | I | GAM | 2000 | 2022 | 18 |
| 32 | Beelbroekstraat | Oost-Vlaanderen | 1.95E-05 | 0.00E+00 | -0.48 | D | GAM | 2013 | 2022 | 10 |
| 41 | Bergstraat (Kemmel) | West-Vlaanderen | 3.42E-01 |  | 0.00 | NS | GAM | 1988 | 2021 | 30 |
| 43 | Bergstraat | Antwerpen | 4.68E-05 | 9.82E-02 | -0.03 | F | GAM | 1993 | 2022 | 25 |
| 45 | Berlaarbaan | Antwerpen | 2.53E-02 | 5.89E-04 | -0.15 | D | GAM | 2007 | 2022 | 10 |
| 60 | Bleukstraat | Antwerpen | 3.14E-04 | 2.81E-10 | -0.26 | D | GAM | 2011 | 2022 | 11 |
| 61 | Blinckaertduinbos | West-Vlaanderen | 3.55E-03 | 6.76E-01 | 0.02 | F | GAM | 2007 | 2022 | 10 |
| 67 | Borggravevijverstraat | Limburg | 7.20E-04 | 2.30E-03 | -0.04 | D | GAM | 1988 | 2017 | 17 |
| 79 | Bramensdam (Bazel) | Oost-Vlaanderen | 6.31E-03 | 2.14E-06 | 0.14 | I | GAM | 2002 | 2022 | 13 |
| 86 | Buitenland | Antwerpen | 1.61E-03 | 2.17E-08 | -0.13 | D | GAM | 2013 | 2022 | 10 |
| 89 | Burchtstraat (Kolmont) | Limburg | 1.73E-02 | 1.03E-01 | -0.02 | F | GAM | 1989 | 2022 | 11 |
| 91 | Bussegem | Oost-Vlaanderen | 3.42E-01 |  | 0.00 | NS | GAM | 2012 | 2022 | 11 |
| 100 | Damstraat (Opdorp) | Oost-Vlaanderen | 9.34E-09 | 0.00E+00 | -0.17 | D | GAM | 2011 | 2022 | 12 |
| 105 | De Pont | Antwerpen | 4.54E-02 | 1.00E+00 | 0.00 | F | GAM | 2006 | 2022 | 17 |
| 106 | De Weert | Antwerpen | 2.77E-03 | 2.37E-09 | -0.13 | D | GAM | 2013 | 2022 | 10 |
| 131 | Eesbeekstraat | Oost-Vlaanderen | 1.59E-01 |  | -0.14 | NS | GAM | 2009 | 2020 | 11 |
| 136 | Eindepoel | Antwerpen | 2.49E-03 | 2.22E-05 | -0.07 | D | GAM | 2012 | 2022 | 12 |
| 138 | Einhovensebaan | Antwerpen | 1.19E-01 |  | -0.04 | NS | GAM | 2004 | 2022 | 15 |
| 152 | Fietsostrade Lint - Lier | Antwerpen | 1.33E-01 |  | -0.06 | NS | GAM | 2013 | 2022 | 10 |
| 156 | Fort 8 | Antwerpen | 4.38E-03 | 1.00E+00 | 0.01 | F | GAM | 2011 | 2022 | 12 |
| 163 | Ganzendam (Vurste) | Oost-Vlaanderen | 2.76E-02 | 4.90E-04 | -0.11 | D | GAM | 2011 | 2022 | 12 |
| 165 | Gaverbosdreef, S. Van De Veldestraat | Oost-Vlaanderen | 2.69E-02 | 9.28E-03 | -0.19 | D | GAM | 2013 | 2022 | 10 |
| 200 | Heiken | Antwerpen | 2.16E-01 |  | 0.00 | NS | GAM | 1986 | 2022 | 23 |
| 202 | Heirbaan | Vlaams Brabant | 7.97E-03 | 1.67E-03 | -0.25 | D | GAM | 2013 | 2022 | 10 |
| 209 | Hobokense Polder | Antwerpen | 2.28E-01 |  | -0.02 | NS | GAM | 1996 | 2022 | 19 |
| 211 | Hoekstraat (Heppen) | Limburg | 4.89E-02 | 4.93E-02 | -0.04 | D | GAM | 2002 | 2022 | 21 |
| 215 | Hof Ter Bollendreef (Liezele) | Antwerpen | 3.46E-03 | 3.23E-04 | -0.08 | D | GAM | 1989 | 2019 | 16 |
| 227 | Honegemstraat | Oost-Vlaanderen | 2.62E-01 |  | -0.05 | NS | GAM | 2012 | 2022 | 10 |
| 232 | Hospitaalstraat (Vlamertinge) | West-Vlaanderen | 2.63E-01 |  | 0.01 | NS | GAM | 1988 | 2022 | 12 |
| 245 | Kasteeldreef, Bloemenlei | Antwerpen | 9.09E-11 | 7.49E-13 | -0.18 | D | GAM | 2010 | 2021 | 12 |
| 247 | Kasteelhoekstraat (Hollebeke) | West-Vlaanderen | 2.47E-02 | 1.97E-02 | -0.05 | D | GAM | 1998 | 2022 | 11 |
| 255 | Kattenbroek | Antwerpen | 9.71E-03 | 1.03E-01 | -0.14 | F | GAM | 2009 | 2021 | 11 |
| 258 | Kerkedreef | Antwerpen | 3.83E-03 | 2.90E-03 | -0.13 | D | GAM | 1993 | 2021 | 12 |
| 272 | Kloosterbeekstraat | Limburg | 4.03E-02 | 2.10E-02 | 0.08 | I | GAM | 2003 | 2017 | 12 |
| 273 | Kluisbaan | Antwerpen | 3.60E-03 | 4.19E-04 | -0.16 | D | GAM | 2009 | 2022 | 11 |
| 280 | Kokerellestraat | Oost-Vlaanderen | 4.41E-04 | 4.91E-07 | -0.15 | D | GAM | 2013 | 2022 | 10 |
| 295 | Kruineikestraat | Vlaams Brabant | 2.61E-01 |  | -0.06 | NS | GAM | 2013 | 2022 | 10 |
| 300 | Kwarikweg | Oost-Vlaanderen | 1.59E-03 | 2.94E-01 | -0.02 | F | GAM | 2003 | 2021 | 13 |
| 318 | Lichtaartseweg | Antwerpen | 2.73E-02 | 2.73E-02 | -0.06 | D | GAM | 1998 | 2022 | 10 |
| 319 | Liedermeersweg | Oost-Vlaanderen | 4.74E-05 | 2.50E-05 | -0.04 | D | GAM | 1991 | 2022 | 26 |
| 331 | Lusthoflaan (Wondelgem) | Oost-Vlaanderen | 6.21E-02 |  | 0.07 | NS | GAM | 2012 | 2022 | 11 |
| 334 | Makkegemstraat (Schelderode) | Oost-Vlaanderen | 4.68E-05 | 4.84E-04 | -0.09 | D | GAM | 2001 | 2022 | 22 |
| 344 | Meierij (Schelderode) | Oost-Vlaanderen | 3.42E-01 |  | -0.02 | NS | GAM | 2011 | 2021 | 11 |
| 346 | Meirestraat | Oost-Vlaanderen | 5.91E-03 | 2.80E-04 | -0.10 | D | GAM | 2009 | 2021 | 12 |
| 347 | Merbeekstraat | Vlaams Brabant | 2.47E-02 | 7.40E-01 | 0.02 | F | GAM | 2007 | 2022 | 15 |
| 348 | Mereldreef | Vlaams Brabant | 2.55E-01 |  | 0.05 | NS | GAM | 2011 | 2020 | 10 |
| 351 | Middelberg | Vlaams Brabant | 1.42E-03 | 1.69E-05 | -0.12 | D | GAM | 2012 | 2022 | 11 |
| 353 | Mikse Baan 1 | Antwerpen | 5.31E-03 | 3.65E-04 | -0.12 | D | GAM | 2011 | 2022 | 12 |
| 354 | Mikse Baan 2 | Antwerpen | 7.63E-07 | 0.00E+00 | -0.24 | D | GAM | 2011 | 2022 | 12 |
| 360 | Molenlei | Antwerpen | 5.85E-02 |  | -0.06 | NS | GAM | 2011 | 2022 | 11 |
| 361 | Molenschansweg | Limburg | 2.49E-03 | 3.30E-01 | -0.01 | F | GAM | 1988 | 2022 | 14 |
| 367 | Moorselestraat | West-Vlaanderen | 1.00E-01 |  | -0.25 | NS | GAM | 2012 | 2021 | 10 |
| 374 | Nachtegaalstraat | West-Vlaanderen | 4.20E-01 |  | -0.02 | NS | GAM | 2003 | 2022 | 18 |
| 384 | Nijlensesteenweg | Antwerpen | 1.59E-01 |  | 0.01 | NS | GAM | 1998 | 2022 | 14 |
| 386 | Normandiestraat | West-Vlaanderen | 5.44E-04 | 2.86E-11 | -0.13 | D | GAM | 2011 | 2022 | 12 |
| 396 | Opstraat | Limburg | 2.49E-03 | 4.26E-04 | 0.07 | I | GAM | 1995 | 2022 | 12 |
| 398 | Oude Galgenstraat | Antwerpen | 1.24E-04 | 8.53E-03 | 0.06 | I | GAM | 2001 | 2021 | 13 |
| 401 | Oude Maasstraat | Limburg | 9.73E-02 |  | -0.05 | NS | GAM | 2001 | 2021 | 12 |
| 403 | Oude Schansstraat (Zelem) | Limburg | 1.44E-02 | 8.13E-03 | -0.03 | D | GAM | 1994 | 2022 | 25 |
| 404 | Oude Scheldestraat (Kaaihoeve) | Oost-Vlaanderen | 9.76E-08 | 0.00E+00 | -0.19 | D | GAM | 2013 | 2022 | 10 |
| 410 | Paddenbroek | Oost-Vlaanderen | 3.36E-02 | 1.66E-02 | -0.04 | D | GAM | 1996 | 2022 | 13 |
| 414 | Palokenstraat | Vlaams Brabant | 3.33E-04 | 2.22E-07 | 0.18 | I | GAM | 2007 | 2022 | 14 |
| 415 | Pannehuisstraat | Limburg | 7.10E-02 |  | -0.02 | NS | GAM | 2001 | 2022 | 16 |
| 418 | Pardasssenhoek | Oost-Vlaanderen | 5.25E-02 |  | -0.12 | NS | GAM | 2011 | 2021 | 10 |
| 426 | Peerlaarstraat | Antwerpen | 1.26E-02 | 1.00E+00 | 0.01 | F | GAM | 2013 | 2022 | 10 |
| 428 | Perreveld | Vlaams Brabant | 3.32E-03 | 6.05E-05 | -0.12 | D | GAM | 2001 | 2022 | 16 |
| 431 | Pierlapont (Loppem) | West-Vlaanderen | 1.19E-08 | 0.00E+00 | -0.23 | D | GAM | 2012 | 2022 | 11 |
| 450 | Reitstraat (Helchteren) | Limburg | 4.68E-05 | 2.72E-08 | -0.09 | D | GAM | 2009 | 2019 | 10 |
| 455 | Reukenstraat | Vlaams Brabant | 1.43E-05 | 0.00E+00 | -0.42 | D | GAM | 2013 | 2022 | 10 |
| 457 | Rhodesgoed (Kachtem) | West-Vlaanderen | 1.11E-06 | 0.00E+00 | -0.22 | D | GAM | 2011 | 2022 | 12 |
| 462 | Rode Dreef | Antwerpen | 5.77E-02 |  | -0.09 | NS | GAM | 2011 | 2022 | 12 |
| 467 | Romeinse Kassei (Voort) | Limburg | 5.38E-06 | 2.53E-15 | -0.16 | D | GAM | 2012 | 2022 | 11 |
| 474 | Rubenskasteel (Weerde) | Vlaams Brabant | 1.08E-01 |  | -0.05 | NS | GAM | 2011 | 2022 | 12 |
| 480 | Schaapstraat | Antwerpen | 4.20E-05 | 4.39E-09 | -0.11 | D | GAM | 2006 | 2022 | 14 |
| 488 | Schilder Evenepoelstraat | Vlaams Brabant | 8.14E-05 | 1.28E-08 | -0.10 | D | GAM | 2007 | 2021 | 11 |
| 489 | Schipdonkbrug | Oost-Vlaanderen | 6.38E-05 | 2.63E-09 | -0.18 | D | GAM | 2008 | 2022 | 14 |
| 492 | Schooldreef | Antwerpen | 4.05E-04 | 5.90E-01 | 0.02 | F | GAM | 2011 | 2022 | 12 |
| 494 | Schoonberg | Oost-Vlaanderen | 3.10E-01 |  | 0.04 | NS | GAM | 2013 | 2022 | 10 |
| 498 | Scouselestraat (Temse) | Oost-Vlaanderen | 2.96E-04 | 3.12E-02 | 0.07 | I | GAM | 2008 | 2022 | 13 |
| 501 | Senthout | Antwerpen | 4.68E-05 | 1.35E-02 | 0.04 | I | GAM | 1994 | 2022 | 23 |
| 516 | Smesstraat | Oost-Vlaanderen | 8.31E-02 |  | 0.09 | NS | GAM | 2011 | 2022 | 11 |
| 518 | Spekstraat (Hallaar) | Antwerpen | 6.19E-01 |  | -0.01 | NS | GAM | 1998 | 2022 | 22 |
| 519 | Spichtstraat | Vlaams Brabant | 2.85E-04 | 8.83E-11 | -0.16 | D | GAM | 2006 | 2022 | 10 |
| 529 | St-Geertruistraat (Neerreppen) | Limburg | 7.46E-03 | 8.09E-03 | -0.08 | D | GAM | 2006 | 2022 | 13 |
| 530 | St-Pietersstraat | West-Vlaanderen | 8.01E-02 |  | -0.05 | NS | GAM | 1998 | 2022 | 11 |
| 532 | Steenberg | Oost-Vlaanderen | 5.77E-02 |  | -0.03 | NS | GAM | 1999 | 2020 | 12 |
| 534 | Steenstortstraat (Beverlo) | Limburg | 1.82E-03 | 7.45E-05 | -0.08 | D | GAM | 2006 | 2021 | 10 |
| 536 | Steenweg Wijchmaal | Limburg | 5.21E-05 | 3.81E-03 | 0.05 | I | GAM | 2004 | 2022 | 18 |
| 542 | Sulferberg (Westouter) | West-Vlaanderen | 2.77E-03 | 1.55E-06 | 0.22 | I | GAM | 1993 | 2003 | 10 |
| 552 | Toekomstlaan | Antwerpen | 4.68E-05 | 4.51E-03 | -0.03 | D | GAM | 1989 | 2019 | 22 |
| 568 | Vijverstraat | Limburg | 2.96E-02 | 9.71E-02 | -0.04 | F | GAM | 2006 | 2022 | 14 |
| 574 | Voordestraat (Humbeek) | Vlaams Brabant | 3.42E-01 |  | -0.01 | NS | GAM | 2010 | 2022 | 13 |
| 576 | Voort | Antwerpen | 2.49E-03 | 5.34E-05 | -0.14 | D | GAM | 2011 | 2022 | 12 |
| 577 | Vosberg | Antwerpen | 4.54E-02 | 6.50E-01 | -0.02 | F | GAM | 2011 | 2022 | 12 |
| 580 | Vroegeinde | Antwerpen | 2.91E-04 | 4.56E-08 | -0.15 | D | GAM | 2011 | 2022 | 12 |
| 585 | Waasmunsterbaan | Oost-Vlaanderen | 9.34E-09 | 1.86E-10 | -0.06 | D | GAM | 1988 | 2022 | 26 |
| 587 | Wallemote/Wolvenhof | West-Vlaanderen | 1.24E-01 |  | 0.05 | NS | GAM | 2011 | 2021 | 11 |
| 597 | Weehaagstraat (Eksaarde) | Oost-Vlaanderen | 5.26E-05 | 0.00E+00 | -0.40 | D | GAM | 2006 | 2017 | 11 |
| 602 | Wijk Monteval | West-Vlaanderen | 3.76E-02 | 1.33E-01 | -0.10 | F | GAM | 2011 | 2022 | 10 |
| 606 | Wilderhof | Vlaams Brabant | 9.09E-11 | 2.53E-15 | -0.32 | D | GAM | 2009 | 2022 | 11 |
| 607 | Wildersedijk | Antwerpen | 6.71E-01 |  | 0.01 | NS | GAM | 1998 | 2022 | 16 |
| 608 | Wilgenbroekstraat | West-Vlaanderen | 2.49E-03 | 3.74E-01 | -0.05 | F | GAM | 1991 | 2003 | 12 |
| 610 | Witte Bomendreef | Vlaams Brabant | 4.11E-01 |  | 0.05 | NS | GAM | 2008 | 2021 | 10 |
| 623 | Zandstraat | Limburg | 3.41E-02 | 2.50E-02 | 0.04 | I | GAM | 1997 | 2022 | 12 |
| 632 | Zeeweg (Sint-Andries) | West-Vlaanderen | 2.16E-01 |  | -0.02 | NS | GAM | 2003 | 2022 | 16 |
| 637 | Zink (Munte) | Oost-Vlaanderen | 2.52E-03 | 5.07E-01 | 0.02 | F | GAM | 2002 | 2021 | 19 |
| 3 | Aan De Rodeberg (Engsbergen) | Limburg |  | 1.75E-01 | 0.07 | NS | GLM | 2017 | 2022 | 6 |
| 6 | Abdijstraat | Vlaams Brabant | | 2.25E-01 | 0.26 | NS | GLM | 2017 | 2021 | 6 |
| 8 | Abstraat (Terlanen) | Vlaams Brabant | | 2.35E-05 | -0.23 | D | GLM | 2014 | 2022 | 9 |
| 20 | Averegten - Boonmarkt (Hallaar | Antwerpen | | 8.15E-01 | -0.02 | NS | GLM | 2015 | 2022 | 8 |
| 23 | Bakvoordestraat | West-Vlaanderen | | 8.72E-01 | -0.01 | NS | GLM | 1990 | 1997 | 6 |
| 31 | Baverstraat (Elderen) | Limburg |  | 6.25E-01 | 0.07 | NS | GLM | 1989 | 1994 | 6 |
| 34 | Beernemsteenweg (Wildenburg) | West-Vlaanderen | | 1.32E-18 | -0.61 | D | GLM | 2013 | 2018 | 6 |
| 38 | Bentstraat | Limburg |  | 4.10E-01 | -0.05 | NS | GLM | 2013 | 2022 | 9 |
| 42 | Bergstraat (Tombeek) | Vlaams Brabant | | 7.33E-02 | -0.21 | NS | GLM | 2014 | 2021 | 8 |
| 47 | Beukendreef | Antwerpen | | 9.66E-01 | 0.00 | NS | GLM | 2014 | 2021 | 6 |
| 52 | Bijlokestraat | Oost-Vlaanderen | | 2.08E-04 | 0.29 | I | GLM | 2013 | 2022 | 9 |
| 64 | Bogaardenstraat | Vlaams Brabant | | 6.26E-01 | -0.01 | NS | GLM | 2008 | 2021 | 9 |
| 70 | Boshoek | Antwerpen | | 5.76E-07 | -0.21 | D | GLM | 1995 | 2002 | 8 |
| 71 | Boskapellaan | Oost-Vlaanderen | | 6.39E-02 | -0.43 | NS | GLM | 2016 | 2022 | 7 |
| 73 | Bosstraat (Koersel) | Limburg |  | 1.69E-01 | -0.13 | NS | GLM | 2015 | 2022 | 8 |
| 74 | Bosstraat | West-Vlaanderen | | 4.47E-01 | -0.17 | NS | GLM | 2013 | 2021 | 8 |
| 77 | Boudewijnlaan En Buurt | West-Vlaanderen | | 8.89E-01 | 0.01 | NS | GLM | 2017 | 2022 | 6 |
| 80 | Brandstraat, Bosmansdreef | Oost-Vlaanderen | | 5.37E-02 | -0.26 | NS | GLM | 2017 | 2022 | 6 |
| 81 | Broekstraat (Blaasveld) | Antwerpen | | 2.37E-02 | -0.04 | D | GLM | 1999 | 2022 | 7 |
| 82 | Broekstraat, Fonteinstraat | Antwerpen | | 3.43E-01 | 0.02 | NS | GLM | 1988 | 1997 | 7 |
| 84 | Bruggenhoek | Oost-Vlaanderen | | 4.58E-01 | 0.11 | NS | GLM | 2013 | 2021 | 6 |
| 96 | Cleynhenslaan | Vlaams Brabant | | 2.22E-02 | -0.35 | D | GLM | 2011 | 2017 | 6 |
| 98 | Daalbroekstraat (Rekem) | Limburg |  | 4.39E-03 | -0.19 | D | GLM | 2013 | 2021 | 9 |
| 101 | De 3 Bruggen | Antwerpen | | 6.66E-01 | 0.15 | NS | GLM | 2017 | 2022 | 6 |
| 102 | De Bunt | Oost-Vlaanderen | | 2.47E-04 | -0.12 | D | GLM | 2006 | 2022 | 9 |
| 103 | De Hoef | Vlaams Brabant | | 2.20E-01 | -0.64 | NS | GLM | 2013 | 2015 | 6 |
| 108 | Diepstraat (Loppem) | West-Vlaanderen | | 2.34E-01 | -0.05 | NS | GLM | 1988 | 2001 | 8 |
| 110 | Dompels | Oost-Vlaanderen | | 6.66E-01 | 0.04 | NS | GLM | 2015 | 2021 | 7 |
| 111 | Donderheide | Antwerpen | | 2.39E-01 | 0.15 | NS | GLM | 2013 | 2022 | 8 |
| 116 | Drengel | Antwerpen | | 4.55E-02 | -0.14 | D | GLM | 2004 | 2022 | 7 |
| 128 | Edmond Ronsestraat (Oostakker) | Oost-Vlaanderen | | 2.26E-01 | 0.28 | NS | GLM | 2013 | 2018 | 6 |
| 130 | Eekhofstraat | West-Vlaanderen | | 6.66E-01 | 0.02 | NS | GLM | 2015 | 2022 | 8 |
| 132 | Eichemstraat (Eichem) | Oost-Vlaanderen | | 3.24E-01 | -0.04 | NS | GLM | 2009 | 2022 | 6 |
| 143 | Engelbamp (Nieuwenhoven) | Limburg |  | 1.19E-06 | -0.05 | NS | GLM | 1988 | 2022 | 6 |
| 145 | Europawijk | Antwerpen | | 8.89E-01 | 0.01 | NS | GLM | 2003 | 2021 | 7 |
| 149 | Ezelstraat | West-Vlaanderen | | 3.63E-02 | -0.16 | D | GLM | 2011 | 2021 | 9 |
| 155 | Fonteinstraat (Blaasveld) | Antwerpen | | 7.28E-01 | -0.02 | NS | GLM | 2007 | 2022 | 6 |
| 158 | Frans Verbeekstraat (Overijse) | Vlaams Brabant | | 4.69E-07 | -0.37 | D | GLM | 2014 | 2022 | 8 |
| 170 | Gistelstraat | West-Vlaanderen | | 1.69E-01 | -0.12 | NS | GLM | 2011 | 2021 | 9 |
| 172 | Goorboslei | Antwerpen | | 6.66E-01 | -0.06 | NS | GLM | 2016 | 2022 | 6 |
| 173 | Goorstraat (Oelegem) | Antwerpen | | 4.12E-01 | -0.03 | NS | GLM | 2012 | 2018 | 6 |
| 176 | Gotegemstraat | Oost-Vlaanderen | | 5.60E-01 | 0.06 | NS | GLM | 2016 | 2022 | 7 |
| 177 | Groenboomgaardstraat | West-Vlaanderen | | 1.38E-05 | -0.33 | D | GLM | 2015 | 2022 | 8 |
| 178 | Groenenhoek | Antwerpen | | 1.19E-06 | -0.21 | D | GLM | 2006 | 2017 | 8 |
| 180 | Groot Westhof (Nieuwkerke) | West-Vlaanderen | | 5.80E-12 | -0.44 | D | GLM | 1981 | 1988 | 6 |
| 187 | Hagaard (Overijse) | Vlaams Brabant | | 3.16E-03 | -0.64 | D | GLM | 2017 | 2022 | 6 |
| 191 | Hauwerzele | Oost-Vlaanderen | | 2.43E-03 | -0.31 | D | GLM | 2014 | 2022 | 9 |
| 193 | Heidestraat (Zemst) | Vlaams Brabant | | 1.58E-01 | -0.10 | NS | GLM | 2017 | 2022 | 6 |
| 195 | Heidestraat - Zuid | Antwerpen | | 1.64E-02 | 0.38 | I | GLM | 2014 | 2021 | 8 |
| 197 | Heidestraat | Vlaams Brabant | | 6.58E-05 | 0.33 | I | GLM | 2011 | 2017 | 7 |
| 198 | Heiken (O.L.V.Waver) | Antwerpen | | 2.20E-01 | -0.08 | NS | GLM | 2014 | 2022 | 8 |
| 210 | Hoek Ter Hulst (Moortsele) | Oost-Vlaanderen | | 3.18E-01 | -0.03 | NS | GLM | 2013 | 2022 | 6 |
| 213 | Hoevendijk | Antwerpen | | 2.52E-02 | -0.10 | D | GLM | 2014 | 2022 | 9 |
| 221 | Hollebeek En Schauselhoekstraa | Oost-Vlaanderen | | 5.23E-03 | -0.25 | D | GLM | 2012 | 2019 | 8 |
| 223 | Holsteenweg | Limburg |  | 9.37E-07 | 0.33 | I | GLM | 2014 | 2022 | 8 |
| 224 | Hommelhofstraat | West-Vlaanderen | | 1.85E-01 | -0.07 | NS | GLM | 2015 | 2022 | 8 |
| 233 | Houtstraat (Olsene) | Oost-Vlaanderen | | 7.81E-01 | 0.05 | NS | GLM | 2013 | 2018 | 6 |
| 234 | Huybergsebaan | Antwerpen | | 3.88E-10 | -0.32 | D | GLM | 1999 | 2018 | 9 |
| 235 | Ijshoutestraat | Oost-Vlaanderen | | 1.69E-10 | -0.67 | D | GLM | 2012 | 2019 | 6 |
| 243 | Karperstraat | West-Vlaanderen | | 4.62E-01 | -0.08 | NS | GLM | 2011 | 2018 | 8 |
| 249 | Kasteelstraat (Dikkelvenne) | Oost-Vlaanderen | | 2.52E-02 | -0.28 | D | GLM | 2011 | 2018 | 7 |
| 261 | Kerkstraat (Tielrode) | Oost-Vlaanderen | | 4.38E-02 | 0.09 | I | GLM | 2008 | 2019 | 9 |
| 266 | Kievitstraat | Antwerpen | | 6.78E-01 | -0.03 | NS | GLM | 2015 | 2022 | 8 |
| 270 | Klare Grachtstraat | West-Vlaanderen | | 1.61E-02 | -0.29 | D | GLM | 2011 | 2019 | 9 |
| 274 | Kluisstraat | Antwerpen | | 1.24E-01 | -0.05 | NS | GLM | 2001 | 2016 | 8 |
| 278 | Knodbaan (Oelegem) | Antwerpen | | 8.89E-01 | 0.01 | NS | GLM | 2011 | 2018 | 7 |
| 285 | Koolskampstraat | West-Vlaanderen | | 1.76E-08 | -0.27 | D | GLM | 2014 | 2021 | 8 |
| 289 | Koutergoedstraat (Oostakker) | Oost-Vlaanderen | | 1.44E-08 | -0.47 | D | GLM | 2016 | 2022 | 6 |
| 290 | Kouterstraat (Overijse) | Vlaams Brabant | | 2.52E-01 | -0.12 | NS | GLM | 2014 | 2022 | 8 |
| 292 | Kraaibornstraat (Lauw) | Limburg |  | 1.01E-01 | -0.19 | NS | GLM | 2014 | 2021 | 8 |
| 296 | Kruiskerkestraat | West-Vlaanderen | | 3.49E-07 | -0.67 | D | GLM | 2011 | 2021 | 7 |
| 299 | Kuikenstraat | Antwerpen | | 6.66E-01 | -0.07 | NS | GLM | 2013 | 2022 | 9 |
| 302 | Lanestraat (Tombeek) | Vlaams Brabant | | 1.95E-08 | -0.15 | D | GLM | 2014 | 2022 | 9 |
| 304 | Lange Maat, Meerstraat | Oost-Vlaanderen | | 6.38E-01 | 0.05 | NS | GLM | 2014 | 2022 | 8 |
| 310 | Legeweg | West-Vlaanderen | | 1.07E-02 | -0.24 | D | GLM | 1990 | 2001 | 8 |
| 311 | Lembergestraat (Landskouter) | Oost-Vlaanderen | | 3.68E-08 | -0.12 | D | GLM | 2011 | 2022 | 8 |
| 322 | Lindebornstraat (Elderen) | Limburg |  | 9.87E-06 | -0.16 | D | GLM | 1989 | 2006 | 9 |
| 326 | Lotenhullestraat | Oost-Vlaanderen | | 4.54E-05 | 0.10 | I | GLM | 2004 | 2022 | 7 |
| 332 | M. Noëstraat | Vlaams Brabant | | 1.63E-08 | 0.17 | I | GLM | 2008 | 2021 | 9 |
| 342 | Meersakkerstraat | Oost-Vlaanderen | | 2.34E-01 | 0.11 | NS | GLM | 2015 | 2022 | 8 |
| 343 | Meersstraat, Scheutlagestraat | Oost-Vlaanderen | | 1.68E-12 | -0.51 | D | GLM | 2015 | 2022 | 8 |
| 356 | Milleniumvijver (Elewijt) | Vlaams Brabant | | 5.16E-02 | 0.12 | NS | GLM | 2012 | 2022 | 9 |
| 375 | Nachtegalenstraat | Vlaams Brabant | | 1.79E-01 | 0.09 | NS | GLM | 2015 | 2021 | 6 |
| 379 | Neremweg | Limburg |  | 1.88E-01 | -0.13 | NS | GLM | 2015 | 2022 | 8 |
| 383 | Nieuwstraat | Vlaams Brabant | | 1.28E-29 | -0.18 | D | GLM | 1991 | 2021 | 7 |
| 385 | Ninoofsesteenweg | Oost-Vlaanderen | | 6.46E-06 | 0.12 | I | GLM | 2012 | 2021 | 9 |
| 399 | Oude Gentweg | Oost-Vlaanderen | | 2.55E-11 | 0.13 | I | GLM | 2003 | 2016 | 8 |
| 400 | Oude Lichterveldsestraat | West-Vlaanderen | | 1.58E-01 | -0.29 | NS | GLM | 2015 | 2020 | 7 |
| 402 | Oude Maria Lindestraat | West-Vlaanderen | | 2.66E-02 | 0.38 | I | GLM | 2013 | 2021 | 7 |
| 421 | Parkstraat | West-Vlaanderen | | 1.45E-09 | -0.63 | D | GLM | 2011 | 2017 | 6 |
| 424 | Pater Penninckxstraat | Vlaams Brabant | | 4.38E-02 | 0.12 | I | GLM | 2015 | 2022 | 8 |
| 440 | Populierenlaan | Vlaams Brabant | | 6.66E-01 | -0.07 | NS | GLM | 2014 | 2021 | 6 |
| 452 | Remerstraat | Vlaams Brabant | | 5.55E-04 | -0.28 | D | GLM | 2011 | 2016 | 6 |
| 453 | Reppelerweg (Grote Brogel) | Limburg |  | 8.54E-02 | -0.10 | NS | GLM | 2005 | 2022 | 7 |
| 454 | Retiesebaan | Antwerpen | | 3.20E-01 | -0.12 | NS | GLM | 2015 | 2022 | 6 |
| 459 | Rijkegemkouter | West-Vlaanderen | | 3.26E-01 | -0.09 | NS | GLM | 2011 | 2021 | 8 |
| 469 | Rondrit In Gemeente | West-Vlaanderen | | 2.57E-01 | -0.20 | NS | GLM | 2013 | 2022 | 6 |
| 471 | Rosendaelweg | Antwerpen | | 4.21E-02 | -0.15 | D | GLM | 2013 | 2022 | 8 |
| 478 | Salphensebaan | Antwerpen | | 9.53E-15 | -0.42 | D | GLM | 2016 | 2022 | 7 |
| 481 | Schalmeidreef | Antwerpen | | 9.66E-01 | 0.00 | NS | GLM | 2008 | 2016 | 7 |
| 486 | Scheldebroeken (Zele-Dijk) | Oost-Vlaanderen | | 9.45E-01 | 0.00 | NS | GLM | 2000 | 2014 | 9 |
| 487 | Scheldeveldstraat | Oost-Vlaanderen | | 6.36E-01 | -0.09 | NS | GLM | 2014 | 2020 | 7 |
| 495 | Schoterheide | Limburg |  | 5.37E-02 | -0.11 | NS | GLM | 2014 | 2022 | 9 |
| 504 | Sint-Annalaan - Heuken | Vlaams Brabant | | 4.77E-02 | 0.04 | I | GLM | 2008 | 2022 | 7 |
| 507 | Sint-Pauluslaan (Huize Walden) | Antwerpen | | 1.18E-01 | -0.12 | NS | GLM | 2014 | 2022 | 9 |
| 521 | Spinele | Oost-Vlaanderen | | 2.01E-01 | -0.14 | NS | GLM | 2014 | 2021 | 7 |
| 526 | Spoorwegstraat | West-Vlaanderen | | 7.53E-01 | -0.05 | NS | GLM | 2013 | 2022 | 9 |
| 535 | Steenstraat (Westende) | West-Vlaanderen | | 5.44E-19 | -0.89 | D | GLM | 2011 | 2018 | 8 |
| 547 | Terluchtestraat (Ruddervoorde) | West-Vlaanderen | | 3.20E-01 | -0.17 | NS | GLM | 1998 | 2003 | 6 |
| 549 | Tillegembos (Sint Michiels) | West-Vlaanderen | | 3.88E-10 | -0.31 | D | GLM | 2006 | 2016 | 7 |
| 550 | Tinnenpotstraat-Lijsterstraat | West-Vlaanderen | | 2.29E-03 | -0.32 | D | GLM | 2013 | 2021 | 9 |
| 553 | Torrestraat (Machelen) | Oost-Vlaanderen | | 7.85E-01 | 0.03 | NS | GLM | 2000 | 2007 | 7 |
| 555 | Tulpenlaan | West-Vlaanderen | | 2.42E-03 | -0.22 | D | GLM | 2015 | 2022 | 8 |
| 558 | Varenstraat | Antwerpen | | 6.27E-01 | 0.12 | NS | GLM | 2014 | 2022 | 7 |
| 559 | Varestraat | Antwerpen | | 1.83E-03 | -0.18 | D | GLM | 1996 | 2004 | 6 |
| 565 | Vijfstraten | Vlaams Brabant | | 8.89E-01 | -0.01 | NS | GLM | 2011 | 2019 | 9 |
| 579 | Vrijbosstraat | West-Vlaanderen | | 6.78E-01 | -0.03 | NS | GLM | 2014 | 2019 | 6 |
| 584 | Waaienburgseweg (Roesbrugge) | West-Vlaanderen | | 1.75E-01 | -0.14 | NS | GLM | 2006 | 2015 | 8 |
| 593 | Waterstraat | Antwerpen | | 8.59E-01 | 0.02 | NS | GLM | 2007 | 2021 | 6 |
| 620 | Zand | Antwerpen | | 1.30E-01 | 0.25 | NS | GLM | 2017 | 2022 | 6 |
| 624 | Zandstraat, Drengel | Antwerpen | | 3.20E-04 | -0.13 | D | GLM | 2012 | 2021 | 9 |
| 626 | Zandstraat (Geel) | Antwerpen | | 7.81E-01 | 0.01 | NS | GLM | 1998 | 2022 | 6 |
| 628 | Zandstraat (Sint-Katelijne-Waver) | Antwerpen | | 3.12E-01 | -0.18 | NS | GLM | 2016 | 2021 | 6 |

Supplementary Table S2:

This Table shows which initial land use components extracted from the BWK-BBK hybrid map make up the final land use components used in the regression model and gives a short description of what they contain.

| Initial Component | Final Component | Description |
| --- | --- | --- |
| Arable land  Agricultural Grass & Shrubs | Agricultural land | Agricultural area, ranging from bare arable fields to meadows and pastures. |
| Woodland | Woodland | Forested area |
| Buildings  Roads  Covered  Uncovered  Grass and Shrubs | Urbanized area | Build-up area and gardens and roadsides, towns, villages, industrial areas,… |
| Water | Water | Bodies of water, ranging from ponds to rivers, lakes and channels. |
| BWK Grasslands | BWK Grasslands | Grasslands indicated on the Biological valuation map that are not classified as agricultural Grass & Shrubs or Water on the BBK. |
| Railways | Railways | Railways |

Supplementary information S3:

The details of this modification can be seen in the following formula, for brevity we assume three different components instead of six:

We know that:

$$X_{1}+X_{2}+X_{3}=1 \leftrightarrow X_{3}=1-(X_{1}+X_{2})$$

So, within a standard linear model this would result in the following equation:

$$Y_{i}=\beta_{0}+\beta_{1}X_{1,i}+\beta_{2}X_{2,i}+\beta_{3}X_{3,i}+\epsilon_{i}$$

$$\leftrightarrow Y_{i}=\beta_{0}+\beta_{1}X_{1,i}+\beta_{2}X_{2,i}+\beta_{3}(1-(X_{2,i}+X_{1,i}))+\epsilon_{i}$$

$$\leftrightarrow Y_{i}={(\beta}_{0}+\beta_{3})+{(\beta}_{1}-\beta_{3})X_{1,i}+(\beta_{2}-\beta_{3})X_{2,i}+ \epsilon_{i}$$

Supplementary Table S4:

The Eigenvalues and the proportion of explained variation of the different PC axes for the PCA ordinations of land use proportions at 100, 500 and 1000 m.

PCA 100 m

|  | PC1 | PC2 | PC3 | PC4 | PC5 | PC6 | PC7 | PC8 | PC9 | PC10 |
| --- | --- | --- | --- | --- | --- | --- | --- | --- | --- | --- |
| Eigenvalue | 2.52 | 1.65 | 1.21 | 1.11 | 1.04 | 0.99 | 0.82 | 0.63 | 0.55 | 0.47 |
| Proportion Explained | 0.23 | 0.15 | 0.11 | 0.10 | 0.095 | 0.090 | 0.074 | 0.057 | 0.050 | 0.043 |
| Cumulative Proportion | 0.23 | 0.38 | 0.49 | 0.59 | 0.68 | 0.77 | 0.85 | 0.91 | 0.96 | 1.00 |

PCA 500 m

|  | PC1 | PC2 | PC3 | PC4 | PC5 | PC6 | PC7 | PC8 | PC9 | PC10 |
| --- | --- | --- | --- | --- | --- | --- | --- | --- | --- | --- |
| Eigenvalue | 4.06 | 2.06 | 1.11 | 1.01 | 0.89 | 0.58 | 0.50 | 0.37 | 0.29 | 0.14 |
| Proportion Explained | 0.37 | 0.19 | 0.10 | 0.092 | 0.081 | 0.052 | 0.045 | 0.034 | 0.027 | 0.013 |
| Cumulative Proportion | 0.37 | 0.56 | 0.66 | 0.75 | 0.83 | 0.88 | 0.93 | 0.96 | 0.99 | 1.00 |

PCA 1000 m

|  | PC1 | PC2 | PC3 | PC4 | PC5 | PC6 | PC7 | PC8 | PC9 | PC10 |
| --- | --- | --- | --- | --- | --- | --- | --- | --- | --- | --- |
| Eigenvalue | 4.48 | 2.09 | 1.09 | 0.98 | 0.84 | 0.52 | 0.40 | 0.33 | 0.20 | 0.080 |
| Proportion Explained | 0.41 | 0.19 | 0.099 | 0.089 | 0.076 | 0.047 | 0.036 | 0.030 | 0.018 | 0.0073 |
| Cumulative Proportion | 0.41 | 0.60 | 0.70 | 0.79 | 0.86 | 0.91 | 0.94 | 0.97 | 0.99 | 1.00 |

Supplementary Fig. S5: PCA ordination biplot for the 100 m and 1000 m buffer scale. Different colours detail different labels of toad population trends. Marginal density graphs describe the density of different populations corresponding to different trend labels.

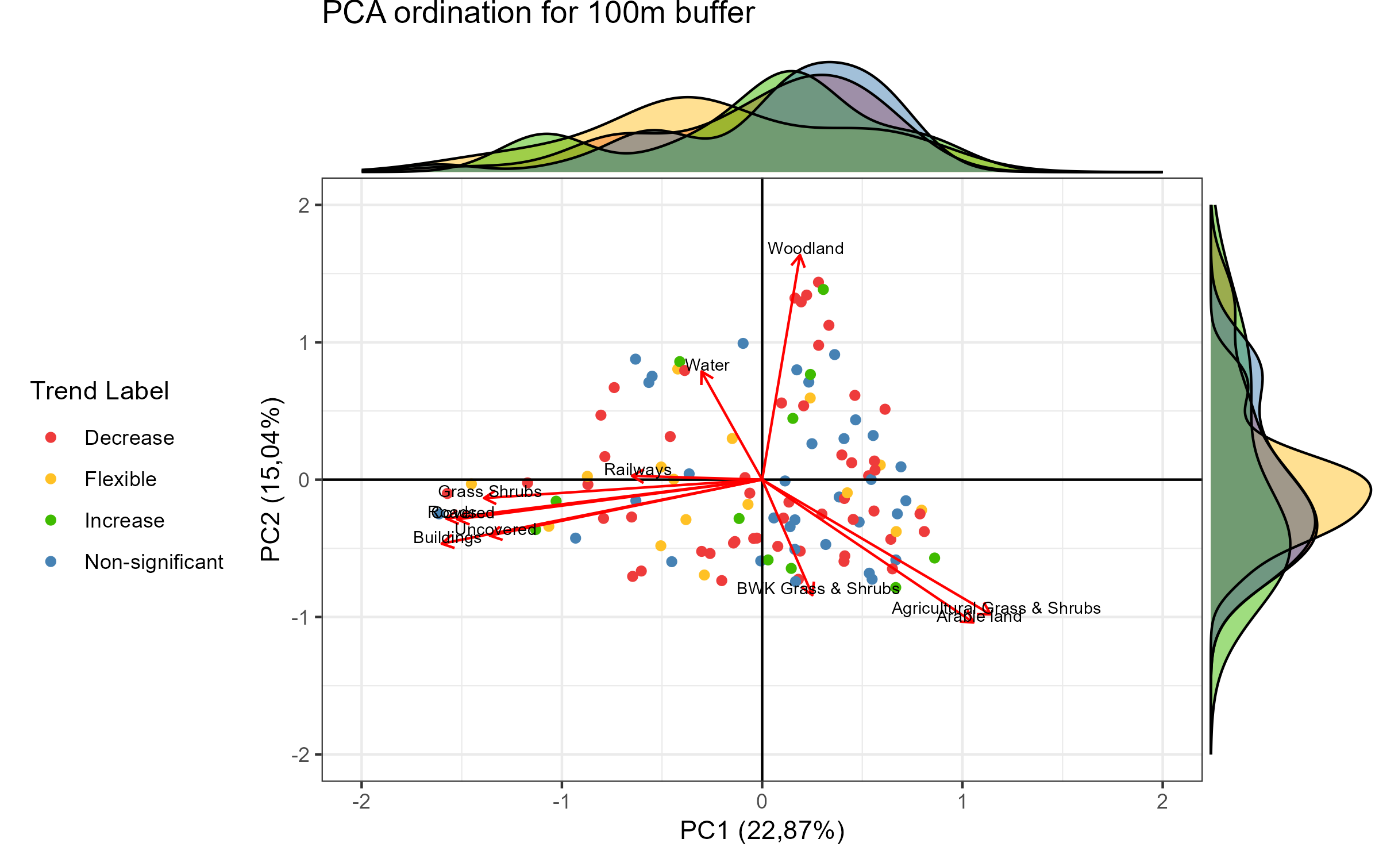

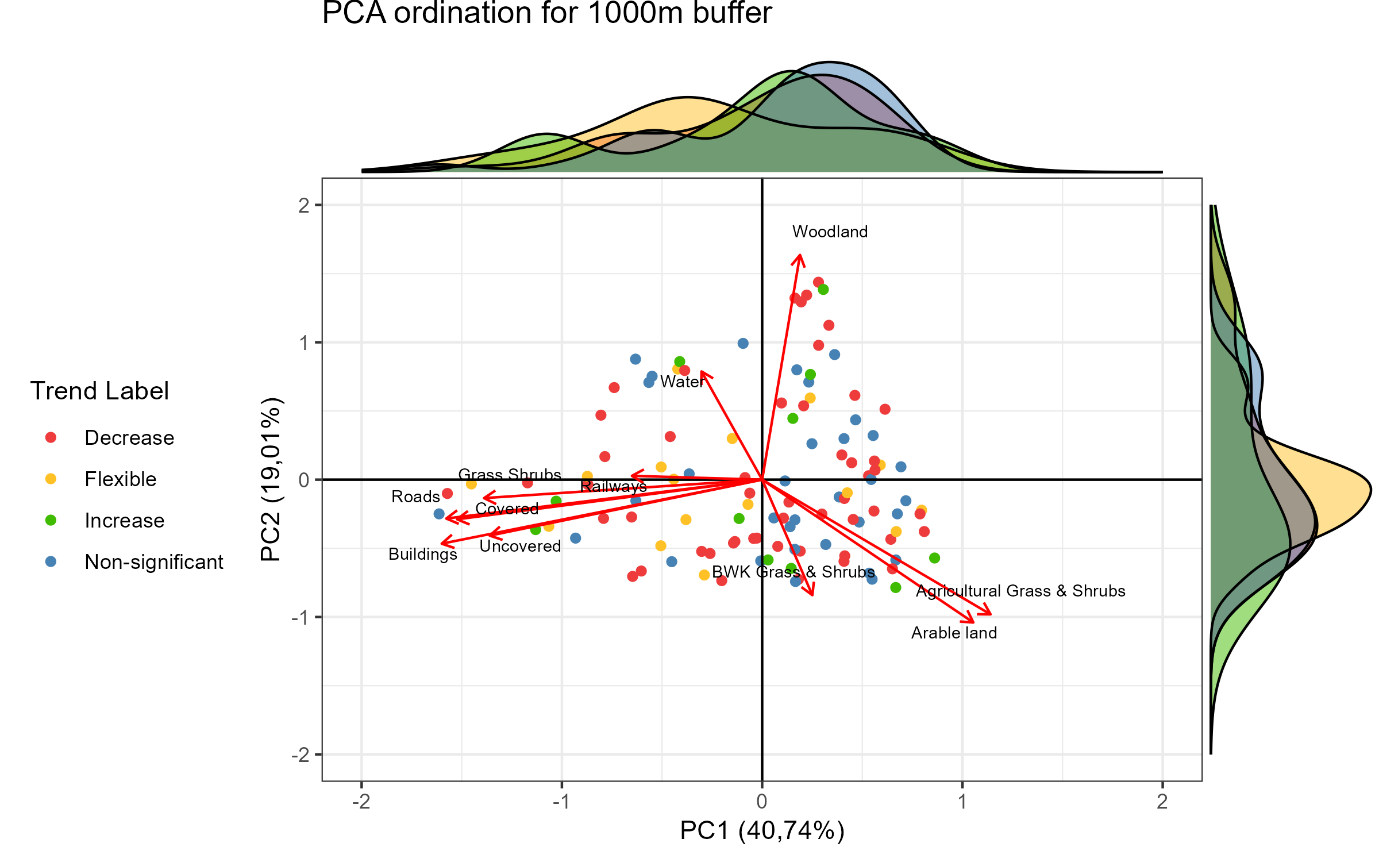
